# Xyloglucan homeostasis and microtubule dynamics synergistically maintain meristem geometry and robustness of phyllotaxis in Arabidopsis

**DOI:** 10.1101/607481

**Authors:** Feng Zhao, Wenqian Chen, Julien Sechet, Marjolaine Martin, Simone Bovio, Claire Lionnet, Yuchen Long, Virginie Battu, Grégory Mouille, Françoise Monéger, Jan Traas

## Abstract

The shoot apical meristem (SAM) gives rise to all aerial organs of the plant. The cell walls are supposed to play a central role in this process, translating molecular regulation into dynamic changes of growth rates and directions, although their precise role in morphogenesis during organ formation remains not well understood. Here we investigate the role of xyloglucans (XyGs), which form a major, yet functionally poorly characterized, wall component in the SAM. Using immunolabeling, biochemical analysis, genetic approaches, micro-indentation, laser ablations and live imaging, we show that XyGs are important for meristem shape and phyllotaxis, although no difference in cell wall stiffness could be observed when XyGs are perturbed. Mutations in enzymes required for XyG synthesis also affect other cell wall components such as cellulose content and the pectin methylation status. Interestingly, we show that the control of cortical microtubules dynamics by the severing enzyme KATANIN becomes vital when XyGs are perturbed or absent. This suggests an active role of the cytoskeleton in compensating for altered wall composition.

## INTRODUCTION

The shoot apical meristem (SAM) gives rise to all aerial organs of the plant. It harbors a pool of stem cells located at the meristem summit, that continuously self-renew and contribute to the formation of new organs (Pfeiffer et al., 2017). These organs are initiated in highly ordered patterns through a process called phyllotaxis. Organ positioning is the result of complex interactions between several hormonal pathway (Galvan-Ampudia et al., 2016). In particular auxin is essential in this process. This hormone accumulates at specific positions through active transport, where it initiates new organs through the activation of a regulatory molecular network (Reinhardt et al., 2003; de Reuille et al., 2006; Rota et al., 2011). How this molecular regulation is then translated into specific growth patterns is not well understood, but it is well established that the cell wall plays a central role.

The cell wall is composed of relatively stiff cellulose microfibrils, embedded in a visco-elastic matrix of polysaccharides (Cosgrove, 2018). In meristematic tissues, cellulose is the most abundant cell wall component, making up 30% of the wall polysaccharides (Yang et al., 2016). The matrix is largely composed of xyloglucans, pectins and arabinans which each make up about 15% of the cell wall (Yang et al., 2016). Work over the last decades has revealed the complexity of wall dynamics and although significant progress has been made, many questions remain concerning the global coordination of wall composition as well as the role of the individual components. The role of cellulose has been relatively well established (Baskin, 2005; McFarlane et al., 2014). The fibrils can be deposited in different arrangements, from completely random to highly aligned arrays. Because of their stiffness, they restrict growth along their length and their orientation largely defines growth directions. Pectins form an important part of the matrix surrounding the cellulose fibrils (Rizk et al., 2000; Cumming et al., 2005). Their precise interaction with other wall components is still not completely understood, but there is strong evidence that pectins participate in regulating organogenesis at the SAM (Peaucelle et al., 2011; Braybrook and Peaucelle, 2013).

Here we focus on the other major matrix component, xyloglucans (XyGs). XyGs are composed of chains of glucose molecules attached through beta (1 -> 4) links, with different sugars as side chains such as xylose, galactose or fucose. They are thought to play a role in both tethering the cellulose microfibrils to other components and in keeping the fibrils separated (Cosgrove, 2018). Their synthesis is controlled by several enzymes. In particular, *XXT1* and *XXT2* encode enzymes with alpha-xylosyltransferase activity that are capable of forming nascent XyG oligosaccharides and their activity is required for XyG synthesis. Another gene, *XYL1*, encodes an alpha-xylosidase that removes the xylose side chains which block the degradation of the backbone (Minic, 2004).

The precise function of XyGs remains controversial. There are several indications that they play an important role at the SAM. Genes encoding xyloglucan endotransglucosylases/hydrolases (XTH), involved in remodeling the XyGs, are very abundantly expressed at the shoot apical meristem (Armezzani et al., 2018). Moreover, Xiao et al (2016) revealed that loss of xyloglucan in the *xxt1/xxt2* mutant affects cell wall integrity, the stability of the microtubule cytoskeleton and the production and patterning of cellulose in primary cell walls in hypocotyls. However, other observations seem to question the role of xyloglucan in morphogenetic events. These include genetic analyses involving mutants of key enzymes required for XyG homeostasis. The *xxt1xxt2* mutant has in the end only a relatively minor growth phenotype compared to what could be expected in the absence of XyGs (Cavalier et al., 2008; Park and Cosgrove, 2012). Likewise, the *xyl1* knock-out mutant showing important modifications in XyG composition (Sampedro et al., 2010; Sampedro et al., 2001; Sechet et al., 2016) is able to form fertile plants.

The SAM, characterized by complex shape changes and growth patterns, offers the possibility to assess wall dynamics and XyG function in a rich developmental context. Using immunolabeling, biochemical analysis and genetic approaches we show that xyloglucans are differentially distributed across the inflorescence meristem, while cellulose and pectins do not appear to exhibit specific distribution patterns. In addition, we have used the *xxt1xxt2* double mutant and the *xyl1* mutant, both perturbed in XyG synthesis as discussed above. The analysis reveals a role for XyG homeostasis in meristem geometry and phyllotaxis. It also points at an active role of the cytoskeleton in compensating for altered wall composition.

## RESULTS

### Xyloglucan distribution patterns correlate with functional domains at the shoot apical meristem

We first examined the distribution of different types of xyloglucans in the wild type SAM using immunolabeling with three different antibodies, LM15, LM25 and LM24. For this purpose, we used both tissue sections and whole mount tissues (respectively Fig. 1A, 1B; Supplemental Fig. S1 for more examples). These antibodies recognize different xyloglucan residues with different affinities (Fig. 1C) (Pedersen et al., 2012).

**Figure 1.**
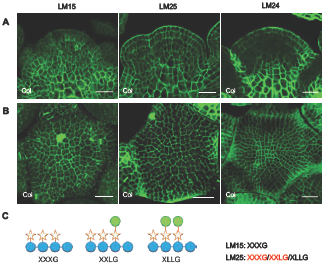
Differential distribution of xyloglucans (XyGs) in Arabidopsis wild-type shoot apices. (A-B) Immunolocalization of XyGs in wild-type (Col) shoot apex sections (A) and whole mount tissues (B) labeled with LM15, LM25 and LM24 antibodies. (C) Schematic structures of XyG subunits and specificity of XyG antibodies. Letters highlighted by red color mean higher affinity. Scale bars, 20µm.

In Col-0 specific patterns were observed:

- The XXXG epitope recognized by LM15, was present throughout the SAM, most strongly in the inner tissues and less in the epidermis and primordia (Fig. 1A). Labelling was particularly striking in differentiating cells at the meristem base, which probably corresponds to the rib meristem. Based on the 3D projection of whole mount signals at the SAM surface, we also found that the XXXG epitope was more abundant in older walls compared to those that had formed more recently (Fig. 1B). This suggests that there is a gradual increase in this epitope throughout the lifespan of the wall.
- The LM25 antibody has a strong affinity for both XXXG and XXLG and a weak affinity for XLLG. Fig. 1A shows a relatively homogeneous labeling across the meristem with this antibody. As indicated above, labeling with LM15 already indicated that XXXG was mainly localized in the internal tissues. Labelling with LM25 would therefore suggest that XXLG is more present in the outer cell layers (Fig. 1A).
- LM24, which mainly detects the XLLG epitope, strongly labels the organ boundaries and the L1 layer, in particular its outer walls and central zone (Fig. 1A, B). LM24 also labels the rib meristem.

In summary, our results on the wild type SAM show specific distribution patterns of XyGs in the SAM, correlated with a number of basic meristem functions, including organ initiation, meristem maintenance and boundary formation.

### Altered XyG content in meristems of *xxt1xxt2* and *xyl1-4* mutants

In order to further investigate the role of XyGs in SAM function, we analyzed *xxt1xxt2* and *xyl1-4*, two mutants affected in enzymes with an opposite effect on XyG side chain branching. As indicated above, whereas XXT1 and XXT2 are responsible for the addition of D-Xylose on the D-Glucose backbone, this D-Xylose residue is removed by XYL1. As shown by *in situ* hybridization, all three genes are expressed at the meristem and show partially overlapping patterns (Supplemental Fig. S2). *XYL1* shows the highest expression in the young initia before the organs start to grow out. As reported by Yang et al (2016), both *XXT1* and *XXT2* are mostly expressed in young primordia (see also supplemental Fig. S2).

We then used the three antibodies mentioned above on the mutants. Although immunolabeling allows only semi-quantitative analysis, we systematically found that, LM15 labeling of XXXG slightly increases throughout the meristems of *xyl1-4* when compared to wild type. (Fig. 1A-B; Fig. 2A-B; Supplemental Fig. S1). LM25 also shows increased labeling throughout the meristem in the mutant (Fig. 1A-B; Fig. 2A-B; Supplemental Fig. S1). LM24 labelling indicates a slight reduction in XLLG mainly in the inner tissues of *xyl1-4* meristems (Fig. 1A; Fig. 2A; Supplemental Fig. S1). Interestingly, *xyl1-4* meristems show a lower signal with LM24 in the central zone of the L1 compared to wild-type (Fig. 1A-B; Fig. 2A-B; Supplemental Fig. S1). As expected, there are no detectable XyGs in *xxt1xxt2* meristems (Fig. 2C; Supplemental Fig. S1).

**Figure 2.**
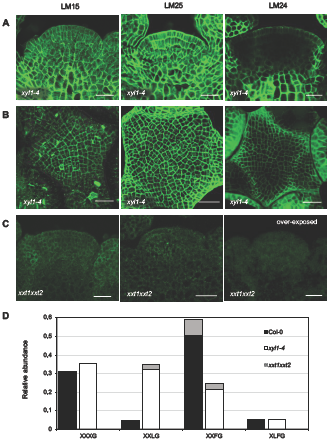
Altered distribution of XyGs in Arabidopsis XyG mutant shoot apices. (A-C) Immunolocalization of XyGs in mutant backgrounds using LM15, LM25 and LM24 antibodies. (A) Sections of *xyl1-4* shoot apices (B), whole mount labelling *of xyl1-4* and (C) sections of *xxt1xxt2* shoot apices. (D) matrix assisted laser-desorption/ ionization time of flight mass spectrometry (MALDI-TOF MS) analysis of XyGs in wild-type, *xyl1-4* and *xxt1xxt2* shoot apices. Grey areas represent the proportion of acetylated subunits. Scale bars, 20µm.

For a more quantitative approach, we carried out an analysis of XyG composition by matrix assisted laser-desorption/ionization time of flight mass spectrometry (MALDI-TOF MS). We dissected 50 meristems of each genotype, which included flower buds younger than stage 3. The results are shown in the Fig. 2D. As expected, we did not find any XyG in *xxt1xxt2* meristems. *xyl1-4* meristems show higher levels of XXLG and a slight increase in XXXG, thus confirming the immunolabeling results. In addition, MALDI-TOF revealed a slight decrease in XLFG residues as well as a reduction in XXFG residues. These changes in XyG fingerprint profiles in the XyG mutants are similar to those found in seedlings (Günl and Pauly, 2011), stems (Sampedro et al., 2010) and embryos (Sechet et al., 2016) suggesting that these enzymes broadly participate in regulating plant development. All together, these results demonstrate that XYL1 and XXT1/2 also regulate XyG composition in the SAM.

### The *xxt1xxt2* mutations affect cellulose content as well as pectin methylation in the meristem

Mutations affecting XyG composition can also lead to alterations of other wall components (Cavalier et al., 2008; Zabotina et al., 2012; Xiao et al., 2016). To test if this was also the case for the SAM, we carried out immunolabeling on wild-type and XyG mutant meristems using a range of antibodies. Pectin and its modifications have been implicated in meristem function (Peaucelle et al., 2008; Peaucelle et al., 2011), but we did not find clear changes in the distribution of methylated (Fig. 3A, B) and de-methylated pectin (Fig. 3 C, D) in XyG mutant meristems. We then further tested the distribution of arabinan, xylan, arabinoxylan and arabinogalactan in XyG deficient mutant meristems and could not find any indication that the distribution of these wall components is perturbed in the mutants (Fig. 3 E-H). Since immunolabeling only provides semi-quantitative information on absolute levels, we performed acid hydrolysis of the cell wall and High Pressure Anion Exchange Chromatography coupled with Pulsed Amperometric Detection (HPAEC-PAD) analysis using WT and XyG mutant inflorescences to obtain whole monosaccharides profiles. The results presented in Fig. 3 J-M show that there are no dramatic changes in the composition of the walls of the *xyl1-4* mutant, in coherence with the immunolabeling data. By contrast, in *xxt1xxt2* a 22% decrease in cellulose content in line with the results of Cavalier et al. (2008) and Xiao et al. (2016) was observed. In addition, a 12% increase in methylated pectin was found (Fig. 3 J-L). Lastly, a prominent drop in fucose (49%, mean value) and xylose (65%, mean value) was observed in *xxt1xxt2* inflorescences which is consistent with the absence of xyloglucans in this mutant (Fig. 3 M). In conclusion, the severe reduction of XyG levels in *xxt1xxt2* significantly alters cellulose levels and pectin methylation, whereas the effects of *xyl1* are more limited.

**Figure 3.**
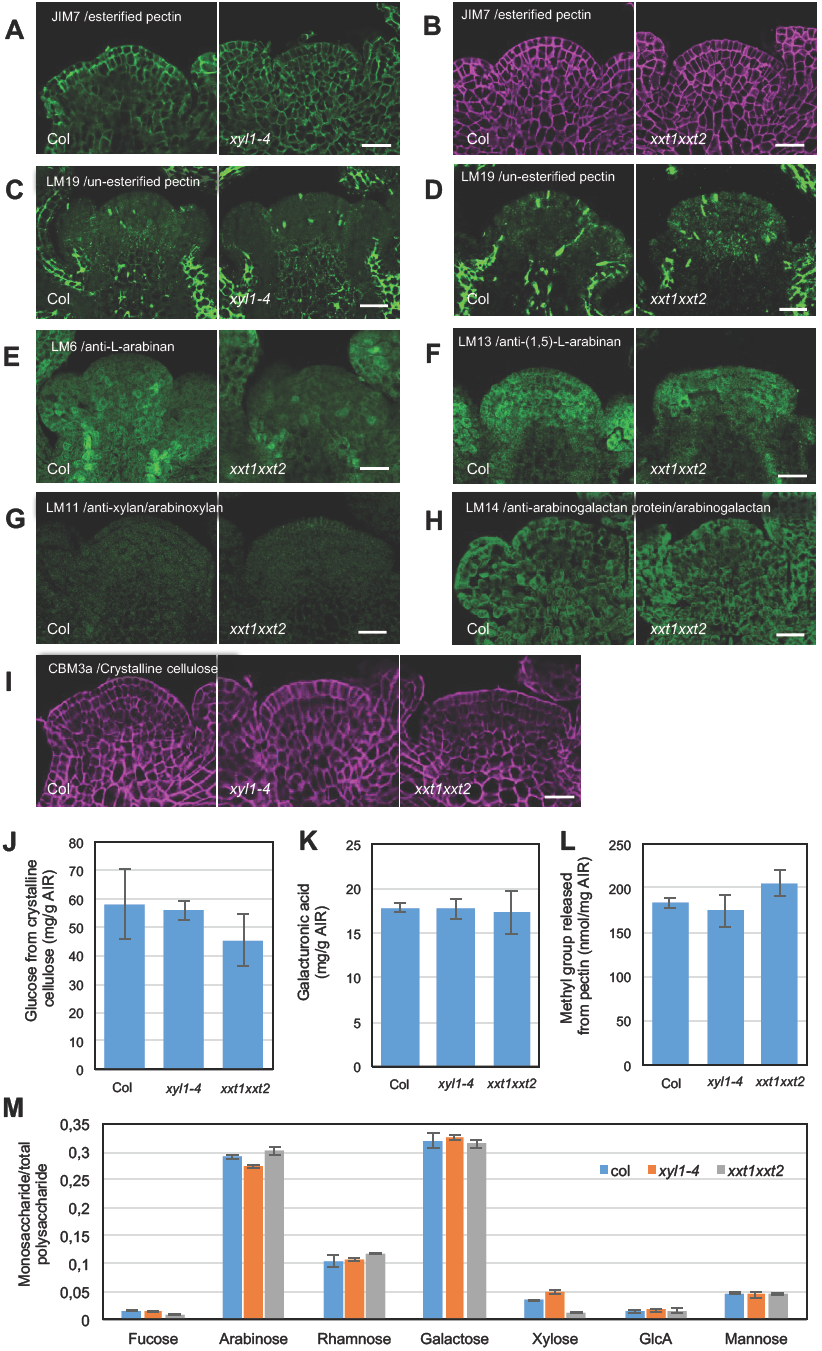
Distribution of other wall components in *xyl1-4* and *xxt1xxt2* SAM. (A-B) Distribution of methylesterified pectin in *xyl1-4* (A) and *xxt1xxt2* (B) SAMs labeled with JIM7 antibody. (C-D) Distribution of un-esterified pectin in *xyl1-4* (C) and *xxt1xxt2* (D) SAMs labeled with LM19 antibody. (E-F) Arabinan distribution pattern in *xxt1xxt2* SAMs labeled with LM6 (E) and LM13 (F) antibodies. (G) Immunolocalization of xylan/arabinoxylan in *xxt1xxt2* SAMs labeled with LM11 antibody. (H) Arabinogalactan distribution pattern in *xxt1xxt2* SAMs labeled with LM14 antibody. (I) Distribution of crystalline cellulose in *xyl1-4* and *xxt1xxt2* SAMs labeled with CBM3a antibody. (J-M) HPAEC-PAD analysis of relative amounts of cellulose (J), homogalacturonan (K), methylesterified pectin (L) and other polysaccharides (M) in XyG mutant inflorescences. AIR, alcohol insoluble residue; scale bars, 30µm.

### Altered XyG composition affects meristem shape and phyllotaxis

We next analyzed the *xyl1* and *xxt1xxt2* phenotypes. The *xyl1-4* mutant has smaller rosette leaves (Supplemental Fig. S3A, B), a phenotype previously also described for *xxt1xxt2* (Park and Cosgrove, 2012; Xiao et al., 2016). Inflorescence stems of both mutants are not growing straight and seem to be partially agravitropic (Supplemental Fig. S3 C and Xiao et al., (2016)). In addition, we observed problems with phyllotaxis in both mutants as shown in Fig. 4A. The *xyl1-4* mutant exhibits a more variable angle distribution when compared to the wild type, with an extra peak at 240° (Fig. 4B-C). Perturbation in phyllotaxis was also observed in *xxt1xxt2* mutants but with different characteristics as the divergence angles in *xxt1xxt2* are often smaller than 137.5° (Fig. 4B, C) and show a peak at 120°. This change in phyllotaxis could possibly be explained by a post meristematic twisting of the cell files along the stem. However we could not detect any evidence for this (Supplemental Fig. S4). It was therefore likely that the changes in phyllotaxis mainly occur at the meristem. To confirm this, we used 3D reconstructions from confocal images and measured successive angles between flower primordia and young flowers on the SAM as described in Fig. 4D. The distribution of divergence angles on SAM is broader in both *xyl1-4* (n= 6 plants and 55 angles) and *xxt1xxt2* (n= 8 plants and 46 angles) compared to Col-0 (n= 10 plants and 67 angles) (Fig. 4E), showing that the organ initiation pattern is perturbed in both mutants. This goes along with changes in meristem shape and size. As shown in the Fig. 5, *xyl1-4* has a flat meristem (Fig. 5A and D) when compared to the wild type. A more detailed quantitative analysis with cellular resolution shows that average cell size is comparable to wild-type in *xyl1-4* (Fig. 5A, B) and therefore is not correlated with these changes in overall geometry (Fig. 5A and D). The meristem of *xxt1xxt2* is flatter and smaller than the wild-type (Fig. 5A, C and D). Cell size is not altered in the mutant (Fig. 5B), showing that reduced meristem size is correlated with reduced cell numbers. In conclusion, the phyllotactic defects observed in both XyG mutants are likely caused by perturbations in organ initiation at the meristematic level and correlate with changes in SAM size and shape.

**Figure 4.**
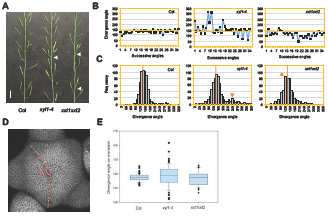
Phyllotactic phenotype of XyG mutants. (A) Representative image showing perturbation of phyllotaxis (indicated by arrow head) in *xyl1-4* and *xxt1xxt2* mutants. (B) Representative distribution angles of siliques on the inflorescence stem of Col, *xyl1-4* and *xxt1xxt2* plants. (C) Distribution of divergence angles of siliques on the Col, *xyl1-4* and *xxt1xxt2* inflorescence stems. Orange lines denote the position of a divergence angle of 137°. Orange arrowheads mark the abnormal angle peaks. n=649 angles from 20 Col plants; n=683 angles from 21 *xyl1-4* plants; n=635 angles from 21 *xxt1xxt2* plants. (D) Diagram showing the method to measure the divergence angles (α) between successive primordia on confocal images of live meristems. (E) Primordia distribution angles on Col, *xyl1-4* and *xxt1xxt2* meristem*s*. 67 angles from 11 Col meristems; 55 angles from 6 *xyl1-4* meristems; 46 angles from 8 *xxt1xxt2* meristems. Scale bar, 1cm in (A).

**Figure 5.**
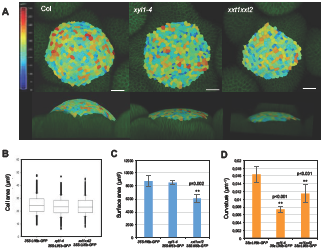
Meristem size and geometry of wild-type and XyG mutants. (A) Overview of meristem size and geometry. Upper panel shows distribution map of cell area on Col and XyG mutant SAMs. Lower panel shows meristem curvature. All plants harbored the plasma membrane marker (*35S:Lti6b-GFP*). Images were post-processed using the MorphoGraphX software. (B) Cell area on meristem surface. n=1409 cells from 4 meristems of *35S:Lti6b-GFP*; n=1469 cells from 4 meristems of *xyl1-4 35S:Lti6b-GFP*; n=1028 cells from 4 meristems of *xxt1xxt2 35S:Lti6b-GFP*. (C) Surface area of Col and XyG mutant meristems calculated from (B). Asterisks denote statistically significant differences from the *35S:Lti6b-GFP* control (student *t*- test, single-tailed). Mean values are represented with SD. (D) Surface curvature of Col and XyGs mutant meristems. n=11 for Col meristems; n=10 for *xyl1-4* meristems and n=8 for *xxt1xxt2* meristems. Asterisks denote statistically significant differences from *35S:Lti6b-GFP* (student *t*-test, single-tailed). Mean values are represented with SD.

### Atomic Force Microscopy (AFM) indentation does not reveal any difference in wall stiffness between mutants and wild type

Changes in geometry and morphogenesis generally result from changes in growth patterns, which in turn largely depend on wall stiffness and the degree of anisotropy. We therefore investigated the mechanical properties of the walls in both mutants using AFM-based nano-indentation on *xyl1-4* and *xxt1xxt2* mutant meristems. We used a silica spherical tip mounted on a silicon cantilever with a nominal force constant of 42 N/m, and a radius of 400 nm (Bovio et al., 2019; see also material and methods). The applied force was of 1 µN, a force corresponding to 100-200 nm indentation, in order to indent the cell wall only (Milani et al., 2011; Tvergaard and Needleman, 2018). Unexpectedly, as shown in Fig. 6, we did not find significant differences between wild type, *xyl1-4* and *xxt1xxt2* at least by applying forces in anticlinal direction on the SAM.

**Figure 6.**
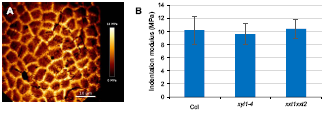
Cell wall elasticity on Col, *xyl1-4* and *xxt1xxt2* SAMs. (A) Representative map of indentation moduli on Col meristem surface. (B) Quantification of Indentation modulus of Col, *xyl1-4* and *xxt1xxt2* SAMs. n=6 for Col meristems, n=7 for *xyl1-4* meristems; n=6 for *xxt1xxt2* meristems. Student *t*-test, single-tailed.

### Microtubule alignment and dynamics are perturbed in XyG mutant meristems

Previous studies have suggested that modified XyG contents can affect cell wall anisotropy and the arrangements of cellulose microfibrils (Xiao et al., 2016). Since microfibril orientations depend on the cortical microtubules (CMTs) guiding the cellulose synthase complexes, we next compared CMT dynamics in wild type and mutants. For this purpose, we introgressed the microtubule reporter construct *p35S:GFP-MBD* into *xyl1-4* and *xxt1xxt2* mutants. Since the GFP-signal was silenced in the *xxt1xxt2* meristem, we used *pPDF1:GFP-MBD* to visualize the microtubules in that mutant. The results are shown in Fig. 7. Confocal imaging revealed that microtubules were less aligned between cells at the meristem in both mutants (Fig. 7A-C, Supplemental Fig. S5 for *in vivo* images), reflecting a reduced coordination of CMTs at the tissue level. These differences were more pronounced in *xyl1-4* and relatively small but significant in *xxt1xxt2*. Interestingly, this seemed to result from different effects at the cellular level. Although the differences are subtle, CMTs are more isotropic in individual cells of *xyl1-4* meristems when compared to the wild type (Fig. 7D), whereas they are more anisotropic in the *xxt1xxt2* mutant meristem (Fig. 7E, Supplemental Fig. S5).

**Figure 7.**
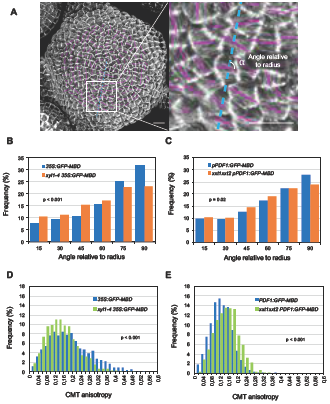
Microtubule patterning on wild-type and XyG mutant SAMs. (A) Representative microtubule patterning on *35S:GFP-MBD* SAM. The orientation and length of magenta bars represent average microtubule orientation and degree of anisotropy in a single cell respectively. Blue lines represent the radius of meristem. Details are enlarged in the right panel. (B-C) Quantifications of CMT orientation relative to the radius of *xyl1-4* (B) and *xxt1xxt2* (C) SAMs. (D-E) Quantifications of CMT anisotropy in *xyl1-4* (D) and *xxt1xxt2* (E) SAMs. Statistic data in (B-E) was calculated from n= 1345 cells of 5 meristems of *35S:GFP-MBD*, n= 1522 cells of 5 meristems of *xyl1-4 35S:GFP-MBD*, n= 1203 cells of 5 meristems of *PDF1:GFP-MBD* and n= 998 cells of 5 meristems of *xxt1xxt2 PDF1: GFP-MBD*. student *t*-test, single-tailed. See also Figure S4 for more details of CMT organization. Scale bar, 10µm in (A).

### Microtubule dynamics may partially compensate the XyG defects in mutant SAMs

There is convincing evidence that CMTs organize in function of mechanical constraints (Hamant et al., 2008; Landrein and Hamant, 2013). The changes in CMT organization observed in the XyG mutants could be due to an altered capacity of the cytoskeleton to reorganize upon mechanical constraints. We tested this capacity in the XyG mutants by performing cell ablations, which cause specific, circumferential rearrangements of the CMT arrays in the cells around the wound. Under our experimental conditions, circumferential microtubule arrays surrounding the wounding start to form two hours after ablation in wild type meristems (Fig. 8A-B). We quantified the microtubule rotation angles after ablation in both XyG mutants and did not find any significant delay in CMT rearrangements when compared to wild type (Fig. 8). These results show that in *xyl1-4* and *xxt1xxt2*, cells have the capacity to perceive exogenous forces and are perfectly able to respond. We therefore hypothesized that the observed changes in CMT anisotropy in the XyG mutants, might be due to an active response of the cytoskeleton to altered wall composition. If this is true, perturbing this response could lead to more severe phenotypes in *xxt1xxt2* or *xyl1-4* backgrounds

**Figure 8.**
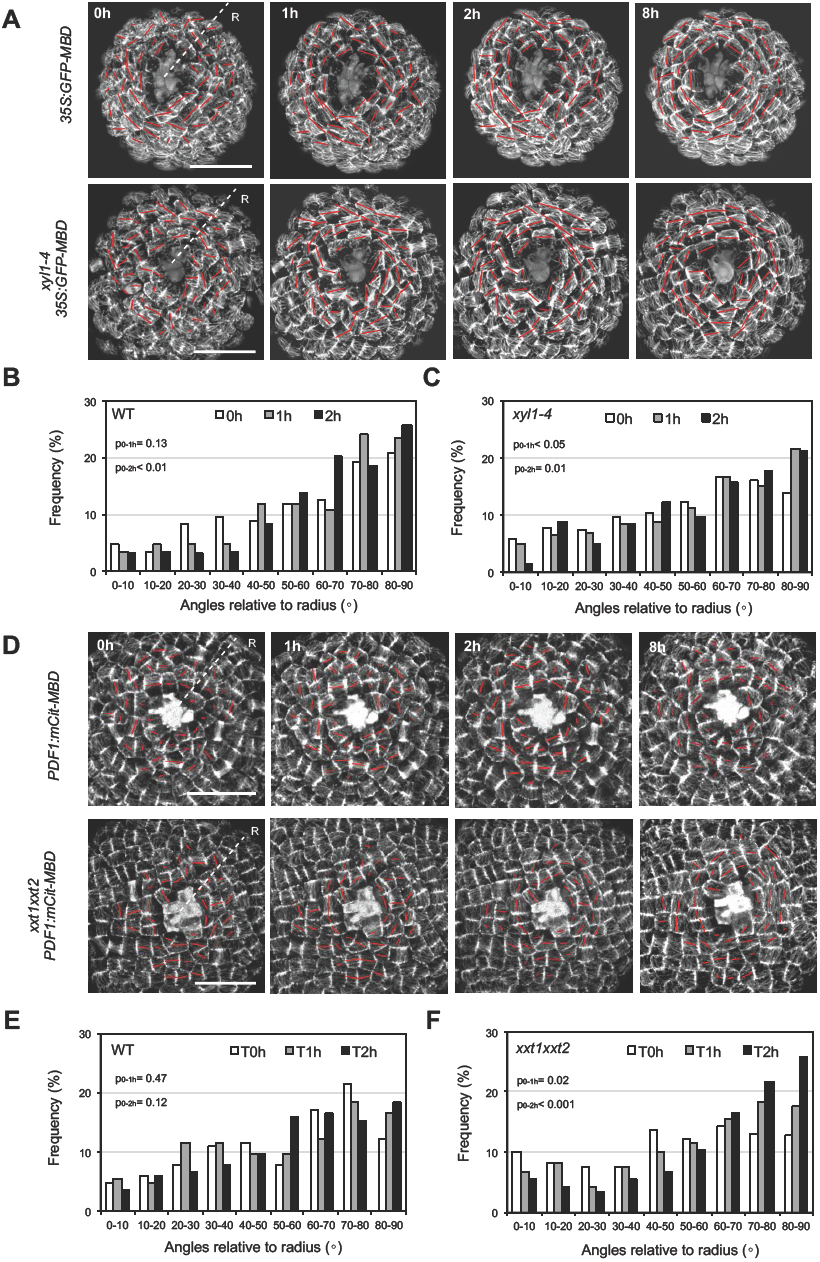
CMT reactions to mechanical perturbation in wild-type, *xyl1-4* and *xxt1xxt2* SAMs. (A) Time series of CMT patterning in *35S:GFP-MBD* (WT) and *xyl1-4 35S:GFP-MBD S*AMs after laser ablation at the meristem center. The orientation and the length of the red bar represent average CMT orientation and degree of CMT anisotropy respectively at cellular level. (B-C) Quantification of CMT orientation angles relative to radius of WT (B) and *xyl1-4* (C) SAMs, 1 hour and 2 hours after laser ablation. n= 167 cells from 4 WT meristems and n=204 cells from 4 *xyl1-4* meristems. (D) Time series of CMT patterning on *pPDF1:mCitrine-MBD* (WT) and *xxt1xxt2 pPDF1:mCitrine-MBD* SAMs after laser ablation at the meristem center. (E-F) Quantification of CMT orientation angles relative to the SAM radius of WT (E), and *xxt1xxt2* (F) SAMs, 1 hour and 2 hours after laser ablation. n= 164 cells from 4 WT meristems and n=181 cells from 4 *xxt1xxt2* meristems. ‘R’ in (A-B) represents radius of meristem; Scale bars, 20µm.

To test this hypothesis we used the *botero* mutant (*bot1/ ktn1*), perturbed in KATANIN, a microtubule severing protein required for microtubule alignment, and introgressed the mutation in *xyl1-4* and *xxt1xxt2*. As shown in Fig. 9, the *xyl1-4 bot1-7* double mutant SAMs have striking concave meristems with a bumpy surface SAM, a phenotype which is much enhanced compared to single mutants (Fig. 9A). In certain individuals, the meristem center was almost hidden between the irregular outgrowths at the surface of the meristem periphery. Several continuous bumps along the orthogonal cutting planes indicated that the coordination of organ growth and separation was affected (Fig. 9B). Consistent with this observation, we observed a dramatic change in phyllotaxis in *xyl1-4bot1-7* double mutant (Fig. 9C-E) compared to WS and *bot1-7* single mutant grown under the same growth condition (Landrein et al., 2015). In view of the irregular surface of the meristems it was sometimes difficult to establish the precise sequence of organ initiation at the meristem in the double mutant. Therefore, we cannot exclude the possibility that the severely perturbed phyllotaxis results from both meristematic and post meristematic events. The cross between *xxt1xxt2* and *ktn1* resulted in even more extreme phenotypes. When we analyzed the offspring of mother plants that were homozygous for *xxt1* and *ktn1* while heterozygous for *xxt2*, we were only able to retrieve four triple mutants in an offspring of 147 plants. These mutants were very small and did not develop beyond the seedling stage (Fig. 10). In conclusion, our results point at negative epistatic interactions, showing that the control of CMT dynamics by KTN becomes vital when XyGs are perturbed or absent.

**Figure 9.**
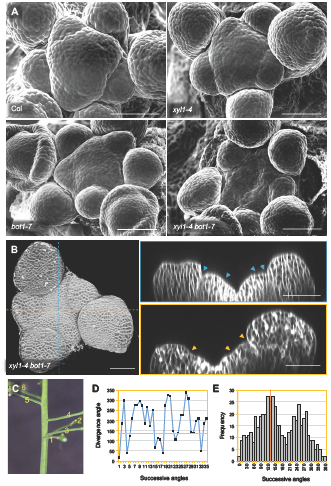
Phenotype of *xyl1-4 bot1-7*. (A) Scanning Electron Microscope (SEM) images of Col, *xyl1-4, bot1-7* and *xyl1-4 bot1-7* SAMs. (B) 3D reconstruction (left panel) and orthogonal view (right panel) of confocal image of *xyl1-4 bot1-7* meristem. The arrowheads mark the points with negative curvatures on meristem surface which are proposed to be organ boundaries. (C) Representative image of silique distribution on *xyl1-4 bot1-7* stem. Numbers denote the silique positions from bottom to top (old to young). (D) Representative silique distribution angles on the inflorescence stem of *xyl1-4 bot1-7*. (E) Distribution of divergence angles of siliques on the *xyl1-4 bot1-7* inflorescence stems. Orange line denotes the position of angle around 137°. n=504 angles from 10 plants. Scale bars=50µm.

**Figure 10:**
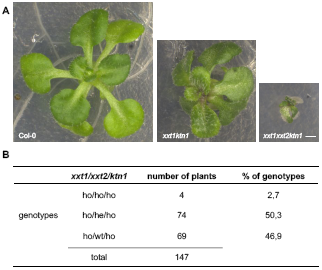
Phenotype of a triple mutant *xxt1xxt2ktn1*. (A) Phenotype of representative Col-0, *xxt1ktn1* and *xxt1xxt2ktn1* plants grown in vitro for 3 weeks. (B) Analysis of the progeny a *xxt1-/- xxt2+/- ktn-/-* plant. Among 147 plants, we identified 4 triple mutants. Scale bar, 1mm.

## DISCUSSION

The precise function of XyGs in development has remained controversial. In a previous study, we showed that genes encoding XyGs modifying enzymes like the XTHs, are highly expressed and show specific expression patterns at the meristem suggesting an important role for XyGs during morphogenesis (Armezzani et al., 2018). Here we explored the role of these components further, and show that specific XyG residues accumulate in different functional domains of the SAM. The organ boundaries and the meristem summit, for example, are characterized by higher LM24 labelling, which probably reflects an increase in XLLG and XXLG subunits. What this precisely implies has yet to be determined, but it should be noted that these domains are characterized by slowly growing cells (Kwiatkowska and Dumais, 2003; Kwiatkowska and Routier-Kierzkowska, 2009). The stiffness of the walls at the boundary has not been studied, but the cells at the meristem summit are slightly more rigid and might be in a particular mechanical, hyperelastic state beyond their linear range of elastic deformation (Kierzkowski et al., 2012; Milani et al., 2014).

The changes in meristem shape are accompanied by modifications in phyllotaxis in both mutants, which can be, at least in part, traced back to early events during organ positioning. There are several possible explanations for this. First, organ outgrowth could be partially impaired, or more irregular, causing young primordia to grow at more variable rates. Such abnormal growth patterns, could destabilize the phyllotactic patterns, for example when an organ grows out more quickly than its predecessor. This type of anomaly, leading to permutations of the positions of successive organs along the stem, has been described for the *ahp6* mutant for example, which is impaired in cytokinin signaling (Besnard et al., 2013). Alternatively, the organ positioning process itself could be modified. As indicated above, organ initiation is caused by the local accumulation of auxin. This accumulation depends on membrane associated auxin transporters of the PIN-FORMED (PIN) family which often show a polar localization. Several studies have pointed at an important role for cell wall components in this polar distribution (Boutté et al., 2006; Heisler et al., 2010; Braybrook and Peaucelle, 2013). Removal of the cell wall during protoplasting, leads to a redistribution of PIN at the cell membrane (Boutté et al., 2006). Certain mutants in cellulose synthase show altered PIN localization in the root (Feraru et al., 2011). It is therefore possible, that the modified cell wall composition in the XyG mutants perturbs phyllotaxis via modified patterns of auxin transport.

The rather mild phenotypes observed after severe changes in such a major wall component remain surprising. Several authors have suggested that this is due to the compensatory action of other cell wall components. In particular pectin has been proposed as a possible source of such a compensation. Although the mutations do not affect the total amount of pectin at the meristem, we did identify a 12% increase in methylesterification in *xxt1xxt2* when compared to wild type, despite the fact that the overall distribution of the different pectin forms was not modified. If we suppose that pectins are the main source for compensation, a 12% increase in pectin methylesterification would then be sufficient. Indeed, the replacement of a carboxyl end with a methyl group changes pectin properties, and potentially affects the dimerization of homogalacturonan mediated by the interaction of Ca^2+^ with un-methylated stretches of galacturonic acid. Note that we did not observe any significant change in wall stiffness, but it remains to be seen if the observed increase in methylesterification would be sufficient to compensate for the absence XyGs.

The results also point at a link between altered XyG content and different aspects of cellulose deposition. The *xxt1xxt2* mutant has a decreased amount of cellulose, whereas the meristems of both *xxt1xxt2* and *xyl1* have modified microtubule dynamics, suggesting altered microfibril deposition as was also reported for hypocotyls (Xiao et al., 2016). In addition the phenotypes are severely enhanced when the XyG mutants are combined with *bot1/ktn1*, impaired in microtubule severing. Therefore, the cytoskeleton seems to compensate at least in part for the loss of XyGs. Indeed, in the absence of XyGs, *bot1/ktn1*, normally able to produce fertile plants, is not able to develop beyond the seedling stage. In addition, combining mutation in *bot1/ktn1* with *xyl1* dramatically increases defects in phyllotaxis and meristem geometry. The altered dynamics of the microtubules most clearly seen in the meristem of *xyl1*, could be due to a reduced or altered capacity of the cytoskeleton to rearrange when cell wall composition is changed as was suggested by Xiao et al (2016). However, the ablation experiments suggest that microtubule dynamics are intact in the mutants. It is therefore reasonable to propose that the altered dynamics of microtubules observed in the mutants reflect some type of active regulation aimed at compensating for the changes in XyG composition. How such a compensation would work is not easy to predict. Part of the answer might come from a role of KTN controlled CMT dynamics in regulating the amount of cellulose, as cellulose levels drop by 20% in stems of the *bot1/ktn1* mutant (Burk et al., 2001). Our own unpublished results even show a 40% drop in the shoot apex. Maintaining the cellulose levels might become critical when XyGs are modified.

The reduced cellulose levels in *bot1/ktn1* raise in turn a number of questions. First, it is not clear why changes in microtubule severing would inhibit the deposition of cellulose so dramatically. Second, cellulose is supposed to contribute significantly to wall stiffness. However, we didn’t measure any important change in the elastic modulus using AFM in *bot1/ktn1* (Uyttewaal et al., 2012; our unpublished results), suggesting that other components compensate for the potential reduction in stiffness due to the loss in cellulose. XyGs are somehow essential in this context, as their presence is absolutely required when KTN is impaired.

In conclusion, XyGs have a significant role in patterning at the shoot apical meristem. This could be due to a direct role of XyG composition in coordinating growth rates and directions, although indirect effects on cell polarity and auxin transport might also be involved. We also find that XyG composition can at least in part compensate for impaired cellulose deposition and vice versa. How this precisely works remains to be elucidated, but the results again illustrate the extraordinary capacity of plant cells to maintain and adapt the properties of their walls to guarantee robust development.

## MATERIALS AND METHODS

### Plant materials and culture conditions

Arabidopsis thaliana Col-0 and Ws-2 ecotype plants were used as wild-type. All mutants and marker lines used in this study have been described previously: *xxt1xxt2*(Cavalier et al., 2008), *xyl1-4* (Sechet et al., 2016), *bot1-7* (Sassi et al., 2014), *ktn1* (Chen et al., 2014), *35S:GFP-MBD* (Hamant et al., 2008), *35S:GFP-Lti6b* (Sassi et al., 2014) *and PDF1:mCitrine-MBD* (Stanislas et al., 2018). The other materials were generated by crossing and subsequently confirmed by genotyping. Plants were grown on soil under long-day condition (16/8 hours LED light period, 150µEm^−2^s^−1^; 60% humidity and 20-22 °C day temperature). For confocal and time-lapse imaging, shoot apices were dissected and cultured *in vitro* on apex culture medium (ACM) as described previously (Sassi et al., 2014).

### Immunolocalization of cell wall components

Inflorescence meristems were infiltrated in the FAA fixative (3.7% formaldehyde, 50% ethanol and 10% acetic acid) under vacuum for 5 min and left in this solution overnight at 4°C. Paraffin embedding sectioning were carried out according to (Zhao et al., 2017). Sections of wild type and mutant specimens were put on the same slide, then deparaffinized in Histo-Clear and rehydrated in a series of ethanol solutions, followed by treatment with membrane permeabilization solution (10% dimethyl sulfoxide (DMSO); 3% nonidet P 40 (NP-40)) for 1 hour. After washing in 1x phosphate buffered saline (PBS) buffer (pH7.0), the slides were incubated with primary antibodies: anti-crystalline cellulose (Plant Probes CBM3a), anti-homogalacturonan (Plant Probes JIM7 and LM19), anti-xyloglucan (Plant Probes LM15, LM24 and LM25), anti-arabinan (Plant Probes LM6 and LM13), anti-xylan/arabinoxylan (Plant Probes LM11), and anti-arabinogalactan (Plant Probes LM14) in 1% (w/v) bovine serum albumin (BSA)/PBS buffer pH7.0 overnight at 4°C, and then the corresponding secondary antibodies: anti-rat IgG (Alexa Fluor 488 conjugated, Molecular Probes A21210 and Dylight 550 Invitrogen SA5-10027) and IgM (Dylight 488 conjugated, Abcam ab96963), anti-goat IgG (Alexa Fluor 488 conjugated, Molecular Probes A11055), anti-mouse IgM (Alexa Fluor 488 conjugated, Invitrogen A-21042) and anti-His tag (Alexa Fluor 555 conjugated, Thermo Fisher MA1-21315-A555) for 3 hours at 37°C. After washing in PBS buffer pH7.0, the slides were observed in a Zeiss LSM 700 laser-scanning confocal microscope. To better detect XyG signals, slides were treated with 0.1% (w/v) pectolyase (Sigma, P5936) in citric acid-sodium phosphate buffer (0.2 M Na_2_HPO_4_, 0.1 M citric acid (pH4.8)) for 45 min before antibody incubation.

For whole mount immunolocalization, dissected shoot apices were fixed in FAA under vacuum for 1 hour. After dehydration and rehydration in a series of ethanol solutions, the shoot apices were digested in a solution containing 0.1% (w/v) pectolyase and 0.1% pectinase (with citric acid-sodium phosphate buffer pH4.8) for 1 hour at room temperature. After membrane permeabilization as described above and washing in 50 mM piperazine-N,N′-bis (PIPES), 5 mM ethylene glycol tetraacetic acid (EGTA), 5 mM MgSO_4_, pH7.0, the shoot apices were incubated with primary anti-xyloglucan antibodies in 3% BSA/0.1% triton/MTBS buffer overnight at 4°C followed by the corresponding secondary antibodies for 3 hours at 37°C. After washing with buffer, the apices were mounted vertically in Murashige and Skoog (MS) medium. Image were taken with a Zeiss LSM700 laser-scanning confocal microscope equipped with water immersion objectives (W N-Achroplan 40x/0.75 M27).

### RNA *in situ* hybridization

RNA in situ hybridization on sections was performed according to (Armezzani et al., 2018) using digoxin-labeled *XYL1* (2924 bp, primers: 5’-ACCATAAGCTAAAGAGGGTTCG and 5’-TAATACGACTCACTATAGGG GAAATGGAGAAGAACAAAACATTACC), *XXT1* (938bp, primers:5’-ATTCTGGGCTAAGCTTCCGTTGand5’-TAATACGACTCACTATAGGG CTCCATACACGACTCCAC) and *XXT2* (538bp, primers: 5’-ATGATTGAGAGGTGTTTAGGAGC and 5’-TAATACGACTCACTATAGGGAGCCATCTCTGCATCGAG) probes from amplified PCR products (prepared according to (Rozier et al., 2014)). Images were taken with Zeiss Axio imager 2 microscope equipped with EC Plan-Neofluar 20x/0.5 objective.

### Cell wall composition analyses

For analyzing the XyG contents on shoot apical meristem (SAM) following the oligosaccharide fingerprinting set up by Lerouxel et al., (2002), 50 shoot apices were dissected and kept in ethanol. After ethanol removal, XyG oligosaccharides were generated by treating samples with endoglucanase in 50 mM sodium acetate buffer, pH 5, overnight at 37°C. Matrix-assisted laser-desorption ionization time of flight mass spectrometry of the XyG oligosaccharides was recorded with a MALDI/TOF Bruker Reflex III using super-DHB (9:1 mixture of 2,5-dihydroxy-benzoic acid and 2-hydroxy-5- methoxy-benzoic acid; Sigma-Aldrich, sigmaaldrich.com) as matrix.

For whole cell wall component measurement, around 0.3g fresh inflorescences were collected for analysis and fixed in 96% ethanol. After grinding in ethanol, they were incubated for 30 min at 70°C. The pellet was then washed twice with 96% ethanol and twice with acetone. The remaining pellet is called alcohol insoluble residues (AIR) and was dried in a fume hood overnight at room temperature.

For pectin measurement, saponification of the AIR (3mg) was performed in triplicates with 0.05 M NaOH. The supernatant containing methyl ester released from the cell wall was then separated from the pellet with polysaccharides. Pectins were extracted from the pellet with 1% ammonium oxalate at 80°C for two hours as described (Krupkova et al., 2007; Mouille et al., 2007; Neumetzler et al., 2012). Galacturonic acid was then quantified by colorimetry using meta-hydroxydiphenyl-sulfuric acid method as described (Blumenkrantz and Asboe-Hansen, 1973). Methyl ester was quantified from NaOH supernatant with a colorimetric assay using enzymatic oxidation of methanol (Klavons and Bennett, 1986).

The monosaccharide composition of the non-cellulosic fraction was determined by hydrolysis of 100 µg AIR with 2 M TFA for 1 h at 120°C. After cooling and centrifugation, the supernatant was dried under a vacuum, resuspended in 200 µl of water and retained for analysis. To obtain the glucose content of the crystalline cellulose fraction, the TFA-insoluble pellet was further hydrolyzed with 72% (v/v) sulfuric acid for 1 h at room temperature. The sulfuric acid was then diluted to 1 M with water and the samples incubated at 100°C for 3 h. All samples were filtered using a 20-µm filter caps, and quantified by HPAEC-PAD on a Dionex ICS-5000 instrument (ThermoFisher Scientific) as described (Sechet et al., 2018).

### Phenotypic analysis (Phyllotaxy, meristem size and geometry measurement)

The phyllotactic patterns were measured as described previously (Besnard et al., 2013). Cell size in the SAMs was obtained by using MorphoGraphX software according to the guideline (https://www.mpipz.mpg.de/MorphoGraphX/help). Only the cells within a domain contained within the organ boundaries were taken into account. Meristem size (surface area) was calculated by summing up all the cell areas per meristem. To calculate meristem curvature, confocal stacks were viewed as two independent orthogonal planes by using orthogonal views function in Fiji software (https://fiji.sc). The radius of meristem was evaluated by drawing a circle tangential to the inner surface of meristem summit. The radius of the circle was taken as the meristem radius (R). The meristem curvature was then calculated as 1/R. To decrease the bias of the measurement, we averaged the curvature value obtained from two orthogonal planes mentioned above on a single meristem. All the data were processed by SigmaPlot and Microsoft Excel software.

### Live imaging and microscopy

For live imaging, dissected meristems were visualized using a membrane marker (*GFP-Lti6b*) under control of an appropriate promoter, or stained with propidium iodide (PI). Samples examined in a Zeiss LSM 700 laser-scanning confocal microscope equipped with water immersion objectives (W Plan-Apochromat 40x/1.0 DIC or W N-Achroplan 40x/0.75 M27). For scanning electron microscopy (SEM), freshly dissected meristems were observed with a HIROX SH-3000 tabletop microscope equipped −20°C and an accelerating voltage of 5kV.

### Image processing and analyses

Fiji software was used for 2D (two dimensional) confocal image analysis. For 3D (three dimensional) image processing, the Zeiss ZEN2 software was used to make a 3D maximum or transparent projection of the signals on meristem. MorphoGraphX software was used to reconstruct the outer meristem surface. To quantify cortical microtubule signals, the images were processed and analyzed according to (Verger et al., 2018) using Fibril tool as described (Boudaoud et al., 2014).

### Atomic Force Microscopy (AFM)

To prevent vibrations, cleanly dissected meristems were fixed vertically on a 60 mm Petri dish (Falcon 60 mm × 15 mm, Corning Ref. 351007) by using biocompatible glue Thin Pour (Reprorubber, Flexbar Ref 16135). AFM experiments were performed on a stand-alone JPK Nanowizard III microscope, driven by a JPK Nanowizard software 6.0. The acquisitions were done using the Quantitative Imaging mode (QI). The experiments have been performed in liquid ACM at room temperature: liquid ACM was added into the petri dish to rehydrate meristems around 1 hour before the beginning of the measurements. We used a silica spherical tip with a nominal radius of 400 nm (Special Development SD-sphere-NCH, Nanosensors) mounted on a silicon cantilever with a nominal force constant of 42 N/m. Scan size was generally of 50 µm with pixel size of 500 nm. The applied force trigger was of 1 µN, a force corresponding to an indentation of 100-200 nm, used in order to indent the cell wall only (Milani et al., 2011; Tvergaard and Needleman, 2018). The ramp size was of 2 µm (1000 data points per curve), approach speed of 100 µm/s and retract speed of 100 µm/s. For more details about cantilever calibration, see Bovio et al., 2019. Data analysis was done using JPK Data Processing software 6.0. Young’s modulus was obtained by fitting the entire force vs tip-sample distance curve with a Hertz model for a sphere. For our analysis, we used a tip radius *R* of 400 nm and a Poisson’s ratio *v* of 0.5 (as it is conventionally set for biological materials), where the Young’s modulus, the point of contact and an offset in force were kept as free parameters of the fit. The same analysis protocol was used on approach and retract curves.

### Laser ablation

We carried out the laser ablation experiments on shoot apical meristems by using a Zeiss LSM 700 laser-scanning confocal microscope, equipped with an Andor MicroPoint (a galvanometer based laser ablation system), which delivered a 6Hz paused laser at 356 nm. Pre-dissected meristems were cultured vertically in ACM for at least four hours prior to the experiment. After being stained with propidium iodide (Sigma, 100mM) for five minutes, the meristems were put under the microscope. Further steps were manipulated by using the iQ software from Andor. Firstly, meristem was visualized to keep focus on the epidermis of meristem summit. Then we used the circular tool in iQ to draw a reign of interest (ROI) with a diameter of 20 pixels (5µm) at the center of the meristem. With a laser power at 8 with 5 repetitions for each point on the ROI, we made a circular wound. To assure the homogeneity of the ablations, the same procedure was carried on all wild-type and XyG mutant meristems.

## ACKNOWLEDGMENTS

We would like to thank Arezki Boudaoud for giving supports on AFM measurement, and Carlos S. Galvan-Ampudia for discussing the project. J.T., F.Z. and W.C. were funded by the ‘Morphodynamics’ ERC grant (NO. 294397). Y.L. was funded by the ERC grant ‘PhyMorph’ (to Arezki Boudaoud). J.S. is supported by a Marie-Curie FP7 COFUND and AgreenSkills+ fellowship (NO. 609398).

## PARTICIPATION

F.Z and W.C. designed and performed experiments and participated in writing. F.M. and J.T. conceived the project and participated in writing the article. J.S and G.M. performed the cell wall analysis. C.L, Y.L. and S.B. participated in imaging experiments, M.M. and V.B. performed genetic analysis.

## LITERATURE CITED

Armezzani A, Abad U, Ali O, Andres Robin A, Vachez L, Larrieu A, Mellerowicz EJ, Taconnat L, Battu V, Stanislas T, et al (2018) Transcriptional induction of cell wall remodelling genes is coupled to microtubule-driven growth isotropy at the shoot apex in *Arabidopsis*. Development 145: dev162255

Baskin TI (2005) Anisotropic expansion of the plant cell wall. Annu Rev Cell Dev Biol 21: 203–222

Besnard F, Refahi Y, Morin V, Marteaux B, Brunoud G, Chambrier P, Rozier F, Mirabet V, Legrand J, Lainé S, et al (2013) Cytokinin signalling inhibitory fields provide robustness to phyllotaxis. Nature 505: 417–421

Blumenkrantz N, Asboe-Hansen G (1973) New method for quantitative determination of uronic acids. Anal biochem 54: 484–9

Boudaoud A, Burian A, Borowska-Wykręt D, Uyttewaal M, Wrzalik R, Kwiatkowska D, Hamant O (2014) FibrilTool, an ImageJ plug-in to quantify fibrillar structures in raw microscopy images. Nat Protoc 9: 457–463

Boutté Y, Crosnier M-T, Carraro N, Traas J, Satiat-Jeunemaitre B (2006) The plasma membrane recycling pathway and cell polarity in plants: studies on PIN proteins. J Cell Sci 119: 1255–1265

Bovio S, Long Y, Monéger F. (2019). Use of Atomic Force Microscopy to measure mechanical properties and turgor pressure of plant cells and plant tissues. JoVE (in press).

Braybrook SA, Peaucelle A (2013) Mechano-chemical aspects of organ formation in Arabidopsis thaliana: the relationship between auxin and pectin. Plos One 8: e57813

Burk DH, Liu B, Zhong R, Morrison WH, Ye Z-H (2001) A katanin-like protein regulates normal cell wall biosynthesis and cell elongation. Plant Cell 13: 807–827

Cavalier DM, Lerouxel O, Neumetzler L, Yamauchi K, Reinecke A, Freshour G, Zabotina OA, Hahn MG, Burgert I, Pauly M, et al (2008) Disrupting two Arabidopsis thaliana xylosyltransferase genes results in plants deficient in xyloglucan, a major primary cell wall component. Plant Cell 20: 1519–1537

Chen X, Grandont L, Li H, Hauschild R, Paque S, Abuzeineh A, Rakusova H, Benkova E, Perrot-Rechenmann C, Friml J (2014) Inhibition of cell expansion by rapid ABP1-mediated auxin effect on microtubules. Nature 516: 90–3

Cosgrove DJ (2018) Diffuse growth of plant cell walls. Plant Physiol 176: 16–27

Cumming CM, Rizkallah HD, McKendrick KA, Abdel-Massih RM, Baydoun EA-H, Brett CT (2005) Biosynthesis and cell-wall deposition of a pectin–xyloglucan complex in pea. Planta 222: 546–555

Feraru E, Feraru MI, Kleine-Vehn J, Martinière A, Mouille G, Vanneste S, Vernhettes S, Runions J, Friml J (2011) PIN polarity maintenance by the cell wall in Arabidopsis. Curr Biol 21: 338–343

Galvan-Ampudia CS, Chaumeret AM, Godin C, Vernoux T (2016) Phyllotaxis: from patterns of organogenesis at the meristem to shoot architecture. Wires Dev Biol 5: 460–473

Günl M, Pauly M (2011) AXY3 encodes a α-xylosidase that impacts the structure and accessibility of the hemicellulose xyloglucan in Arabidopsis plant cell walls. Planta 233: 707–719

Hamant O, Heisler MG, Jonsson H, Krupinski P, Uyttewaal M, Bokov P, Corson F, Sahlin P, Boudaoud A, Meyerowitz EM, et al (2008) Developmental patterning by mechanical signals in Arabidopsis. Science 322: 1650–1655

Heisler MG, Hamant O, Krupinski P, Uyttewaal M, Ohno C, Jönsson H, Traas J, Meyerowitz EM (2010) Alignment between PIN1 polarity and microtubule orientation in the shoot apical meristem reveals a tight coupling between morphogenesis and auxin transport. Plos Biol 8: e1000516

Kierzkowski D, Nakayama N, Routier-Kierzkowska A-L, Weber A, Bayer E, Schorderet M, Reinhardt D, Kuhlemeier C, Smith RS (2012) Elastic domains regulate growth and organogenesis in the plant shoot apical meristem. Science 335: 1096–1099

Klavons JA, Bennett RD (1986) Determination of methanol using alcohol oxidase and its application to methyl ester content of pectins. J Agr Food Chem 34: 597–599

Krupkova E, Immerzeel P, Pauly M, Schmulling T (2007) The TUMOROUS SHOOT DEVELOPMENT2 gene of Arabidopsis encoding a putative methyltransferase is required for cell adhesion and co-ordinated plant development. Plant J 50: 735–50

Kwiatkowska D, Dumais J (2003) Growth and morphogenesis at the vegetative shoot apex of Anagallis arvensis L. J Exp Bot 54: 1585–1595

Kwiatkowska D, Routier-Kierzkowska A-L (2009) Morphogenesis at the inflorescence shoot apex of Anagallis arvensis: surface geometry and growth in comparison with the vegetative shoot. J Exp Bot 60: 3407–3418

Landrein B, Hamant O (2013) How mechanical stress controls microtubule behavior and morphogenesis in plants: history, experiments and revisited theories. Plant J 75: 324–338

Landrein B, Refahi Y, Besnard F, Hervieux N, Mirabet V, Boudaoud A, Vernoux T, Hamant O (2015) Meristem size contributes to the robustness of phyllotaxis in Arabidopsis. J Exp Bot 66: 1317–1324

Lerouxel O, Choo TS, Seveno M, Usadel B, Faye L, Lerouge P, Pauly M (2002) Rapid structural phenotyping of plant cell wall mutants by enzymatic oligosaccharide fingerprinting. Plant Physiol 130: 1754–63

McFarlane HE, Döring A, Persson S (2014) The Cell Biology of Cellulose Synthesis. Annu Rev Plant Biol 65: 69–94

Milani P, Gholamirad M, Traas J, Arnéodo A, Boudaoud A, Argoul F, Hamant O (2011) In vivo analysis of local wall stiffness at the shoot apical meristem in Arabidopsis using atomic force microscopy. Plant J 67: 1116–1123

Milani P, Mirabet V, Cellier C, Rozier F, Hamant O, Das P, Boudaoud A (2014) Matching patterns of gene expression to mechanical stiffness at cell resolution through quantitative tandem epifluorescence and nanoindentation. Plant Physiol 165: 1399–1408

Minic Z, Rihouey C, Do CT, Lerouge P, Jouanin L (2004) Purification and characterization of enzymes exhibiting β-d-xylosidase activities in stem tissues of Arabidopsis. Plant Physiol 135: 867–878

Mouille G, Ralet MC, Cavelier C, Eland C, Effroy D, Hematy K, McCartney L, Truong HN, Gaudon V, Thibault JF, et al (2007) Homogalacturonan synthesis in Arabidopsis thaliana requires a golgi-localized protein with a putative methyltransferase domain. Plant J 50: 605–14

Neumetzler L, Humphrey T, Lumba S, Snyder S, Yeats TH, Usadel B, Vasilevski A, Patel J, Rose JK, Persson S, et al (2012) The FRIABLE1 gene product affects cell adhesion in Arabidopsis. Plos One 7: e42914

Park YB, Cosgrove DJ (2012) Changes in cell wall biomechanical properties in the xyloglucan-deficient xxt1/xxt2 mutant of Arabidopsis. Plant Physiol 158: 465–475

Peaucelle A, Braybrook SA, Le Guillou L, Bron E, Kuhlemeier C, Höfte H (2011) Pectin-induced changes in cell wall mechanics underlie organ initiation in Arabidopsis. Curr Biol 21: 1720–1726

Peaucelle A, Louvet R, Johansen JN, Höfte H, Laufs P, Pelloux J, Mouille G (2008) Arabidopsis phyllotaxis is controlled by the methyl-esterification status of cell-wall pectins. Curr Biol 18: 1943–1948

Pedersen HL, Fangel JU, McCleary B, Ruzanski C, Rydahl MG, Ralet M-C, Farkas V, von Schantz L, Marcus SE, Andersen MCF, et al (2012) Versatile high resolution oligosaccharide microarrays for plant glycobiology and cell wall research. J Biol Chem 287: 39429–39438

Pfeiffer A, Wenzl C, Lohmann JU (2017) Beyond flexibility: controlling stem cells in an ever changing environment. Curr Opin Plant Biol 35: 117–123

Reinhardt D, Pesce E-R, Stieger P, Mandel T, others (2003) Regulation of phyllotaxis by polar auxin transport. Nature 426: 255

de Reuille PB, Bohn-Courseau I, Ljung K, Morin H, Carraro N, Godin C, Traas J (2006) Computer simulations reveal properties of the cell-cell signaling network at the shoot apex in Arabidopsis. Proc Natl Acad Sci USA 103: 1627–1632

Rizk SE, Abdel-Massih RM, Baydoun EA-H, Brett CT (2000) Protein-and pH-dependent binding of nascent pectin and glucuronoarabinoxylan to xyloglucan in pea. Planta 211: 423–429

Rota CL, Chopard J, Das P, Paindavoine S, Rozier F, Farcot E, Godin C, Traas J, Monéger F (2011) A data-driven integrative model of sepal primordium polarity in Arabidopsis. Plant Cell 23: 4318–4333

Routier-Kierzkowska A-L, Weber A, Kochova P, Felekis D, Nelson BJ, Kuhlemeier C, Smith RS (2012) Cellular force microscopy for in vivo measurements of plant tissue mechanics. Plant Physiol 158: 1514–1522

Rozier F, Mirabet V, Vernoux T, Das P (2014) Analysis of 3D gene expression patterns in plants using whole-mount RNA in situ hybridization. Nat Protoc 9: 2464–2475

Sampedro J, Pardo B, Gianzo C, Guitian E, Revilla G, Zarra I (2010) Lack of α-xylosidase activity in Arabidopsis alters xyloglucan composition and results in growth defects. Plant Physiol 154: 1105–1115

Sampedro J, Sieiro C, Revilla G, González-Villa T, Zarra I (2001) Cloning and expression pattern of a gene encoding an α-xylosidase active against xyloglucan oligosaccharides from Arabidopsis. Plant Physiol 126: 910–920

Sassi M, Ali O, Boudon F, Cloarec G, Abad U, Cellier C, Chen X, Gilles B, Milani P, Friml J, et al (2014) An auxin-mediated shift toward growth isotropy promotes organ formation at the shoot meristem in Arabidopsis. Curr Biol 24: 2335–2342

Schillers H, Rianna C, Schape J, Luque T, Doschke H, Walte M, Uriarte JJ, Campillo N, Michanetzis GPA, Bobrowska J, et al (2017) Standardized nanomechanical atomic force microscopy procedure (SNAP) for measuring soft and biological samples. Sci Rep 7: 5117

Sechet J, Frey A, Effroy-Cuzzi D, Berger A, Perreau F, Cueff G, Charif D, Rajjou L, Mouille G, North HM, et al (2016) Xyloglucan metabolism differentially impacts the cell wall characteristics of the endosperm and embryo during Arabidopsis seed germination. Plant Physiol 170: 1367–80

Sechet J, Htwe S, Urbanowicz B, Agyeman A, Feng W, Ishikawa T, Colomes M, Kumar KS, Kawai-Yamada M, Dinneny JR, et al (2018) Suppression of Arabidopsis GGLT1 affects growth by reducing the L-galactose content and borate cross-linking of rhamnogalacturonan-II. Plant J 96: 1036–1050

Sneddon IN (1965) The relation between load and penetration in the axisymmetric boussinesq problem for a punch of arbitrary profile. Int J Eng Sci 3: 47–57

Stanislas T, Platre MP, Liu M, Rambaud-Lavigne LES, Jaillais Y, Hamant O (2018) A phosphoinositide map at the shoot apical meristem in Arabidopsis thaliana. BMC Biol 16: 20

Tvergaard V, Needleman A (2018) Effect of properties and turgor pressure on the indentation response of plant cells. J Appl Mech 85: 061007–061007–8

Uyttewaal M, Burian A, Alim K, Landrein B, Borowska-Wykręt D, Dedieu A, Peaucelle A, Ludynia M, Traas J, Boudaoud A, et al (2012) Mechanical stress acts via katanin to amplify differences in growth rate between adjacent cells in Arabidopsis. Cell 149: 439–451

Verger S, Long Y, Boudaoud A, Hamant O (2018) A tension-adhesion feedback loop in plant epidermis. eLife 7: e34460

Xiao C, Zhang T, Zheng Y, Cosgrove DJ, Anderson CT (2016) Xyloglucan deficiency disrupts microtubule stability and cellulose biosynthesis in Arabidopsis, altering cell growth and morphogenesis. Plant Physiol 170: 234–249

Yang W, Schuster C, Beahan CT, Charoensawan V, Peaucelle A, Bacic A, Doblin MS, Wightman R, Meyerowitz EM (2016) Regulation of meristem morphogenesis by cell wall synthases in Arabidopsis. Curr Biol 26: 1404–1415

Zabotina OA, Avci U, Cavalier D, Pattathil S, Chou Y-H, Eberhard S, Danhof L, Keegstra K, Hahn MG (2012) Mutations in multiple XXT genes of Arabidopsis reveal the complexity of xyloglucan biosynthesis. Plant Physiol 159: 1367–1384

Zhao F, Zheng Y-F, Zeng T, Sun R, Yang J-Y, Li Y, Ren D-T, Ma H, Xu Z-H, Bai S-N (2017) Phosphorylation of SPOROCYTELESS/NOZZLE by the MPK3/6 kinase is required for anther development. Plant Physiol 173: 2265–2277

