## Supplemental Figures for "Xyloglucan homeostasis and microtubule dynamics synergistically maintain meristem geometry and robustness of phyllotaxis in Arabidopsis"

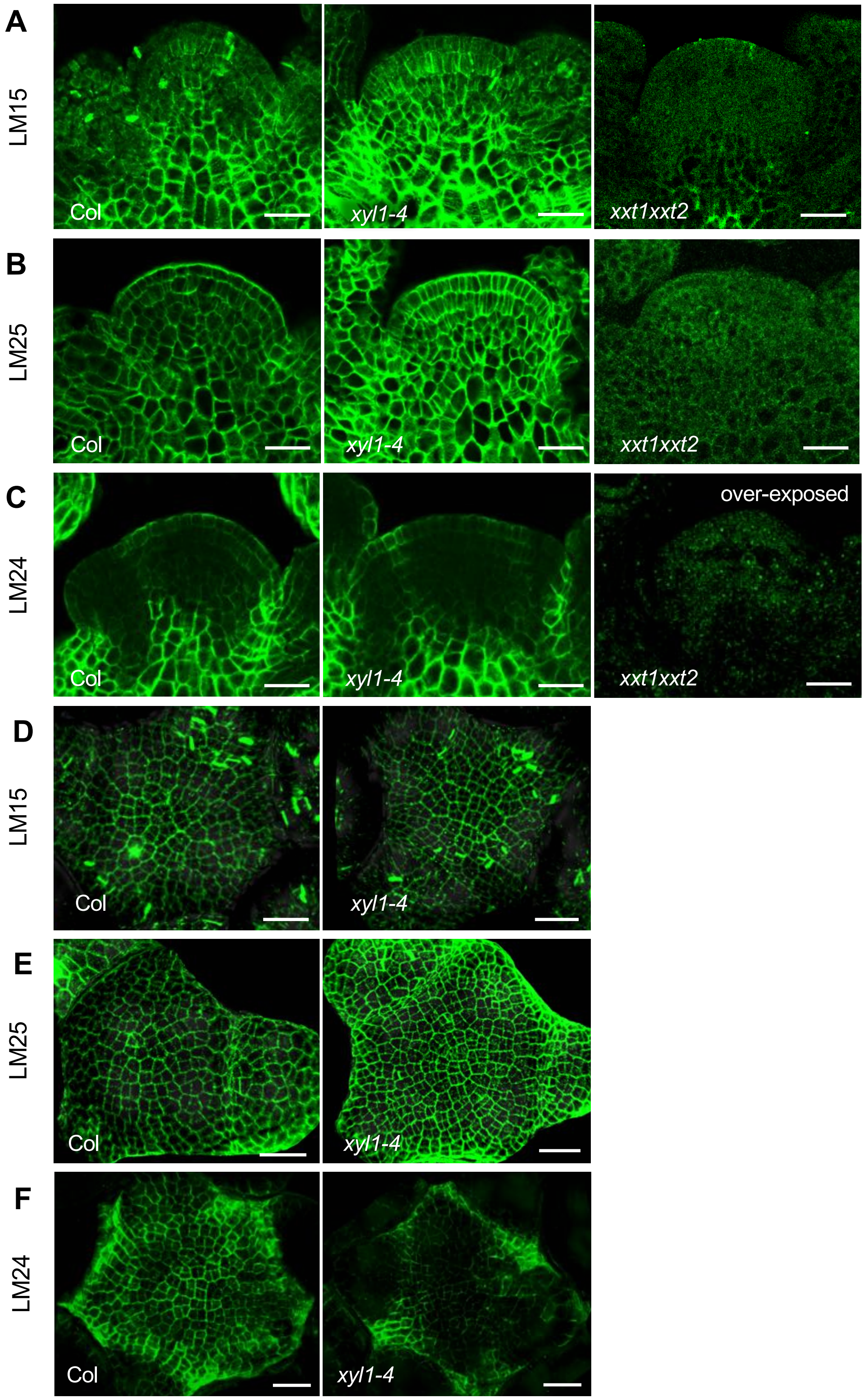


**Figure S1. XyG distribution patterns in wild-type (Col) and XyG mutant shoot apices.** Immunolocalization of XyGs in sections of Col and XyG mutant apices (A-C) and whole mount tissues (D-F) labeled with LM15, LM25 and LM24 antibody. Scale bars, 20μm.

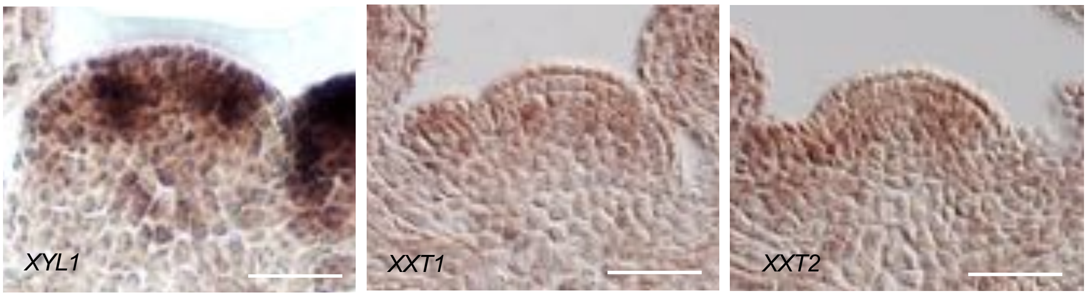


**Figure S2. In situ hybridization of *XYL1*, *XXT1* and *XXT2* in wild-type shoot apices.**

Scale bars, 50μm.

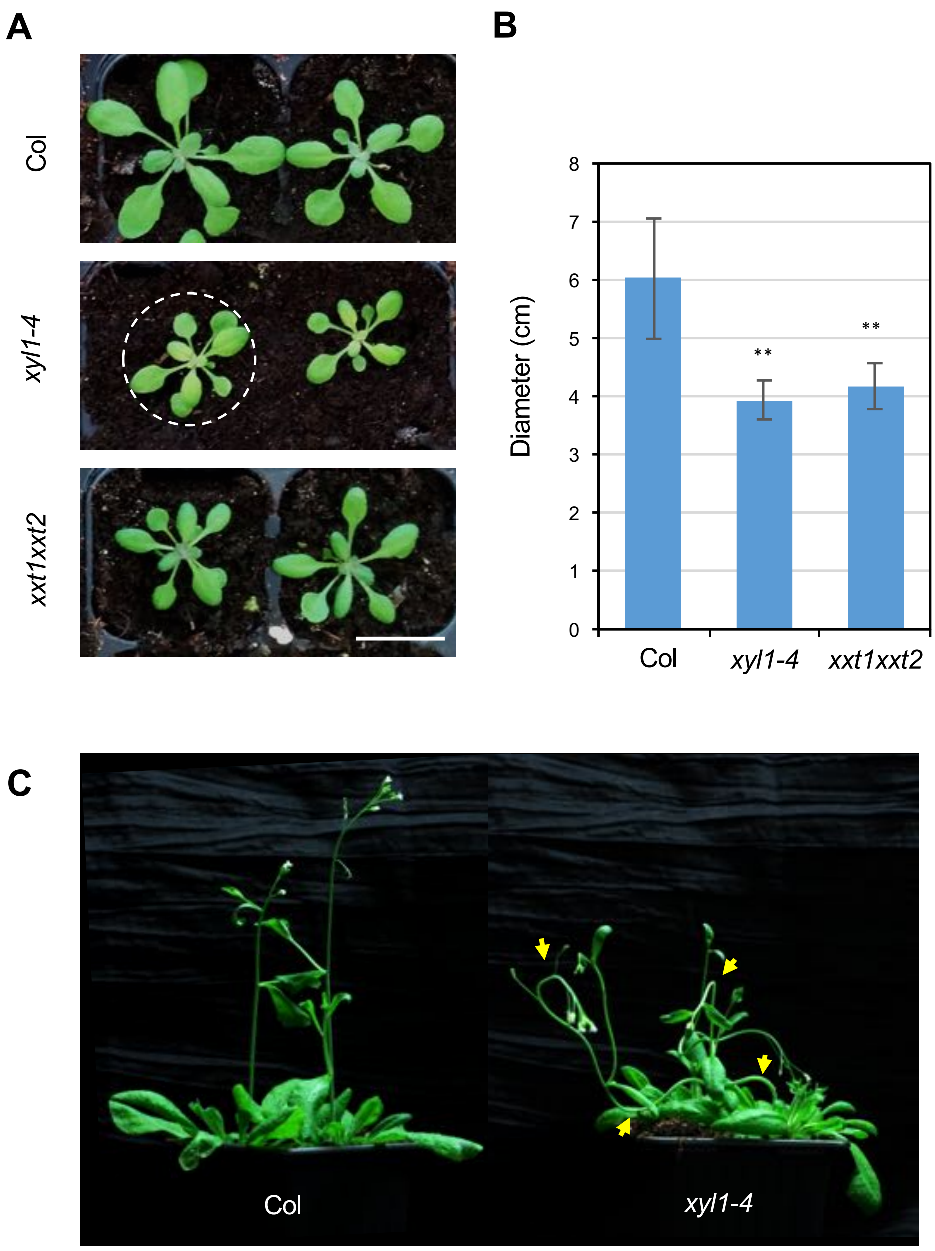


**Figure S3. Phenotype of xyloglucan mutants.**

(A) wild-type (Col) and XyG mutant plants 20DAS (days after stratification). Dashed circle represents the circumscribed circle of the plantlet. This circle is used to quantify the diameter of the plantlet.

(B) Quantification of diameter of plantlets of wild-type (n=12), *xyl1-4* (n=12) and *xxt1xxt2* (n=12). Asterisks denote statistically significant differences (** $p<0.001$; student *t*-test, one-tailed). Mean values are shown with SD.

(C) The gravitropism defects in *xyl1-4*. Yellow arrow-heads indicate the curving stems. The plants are at 35DAS. Scale bar, 2cm in (A).

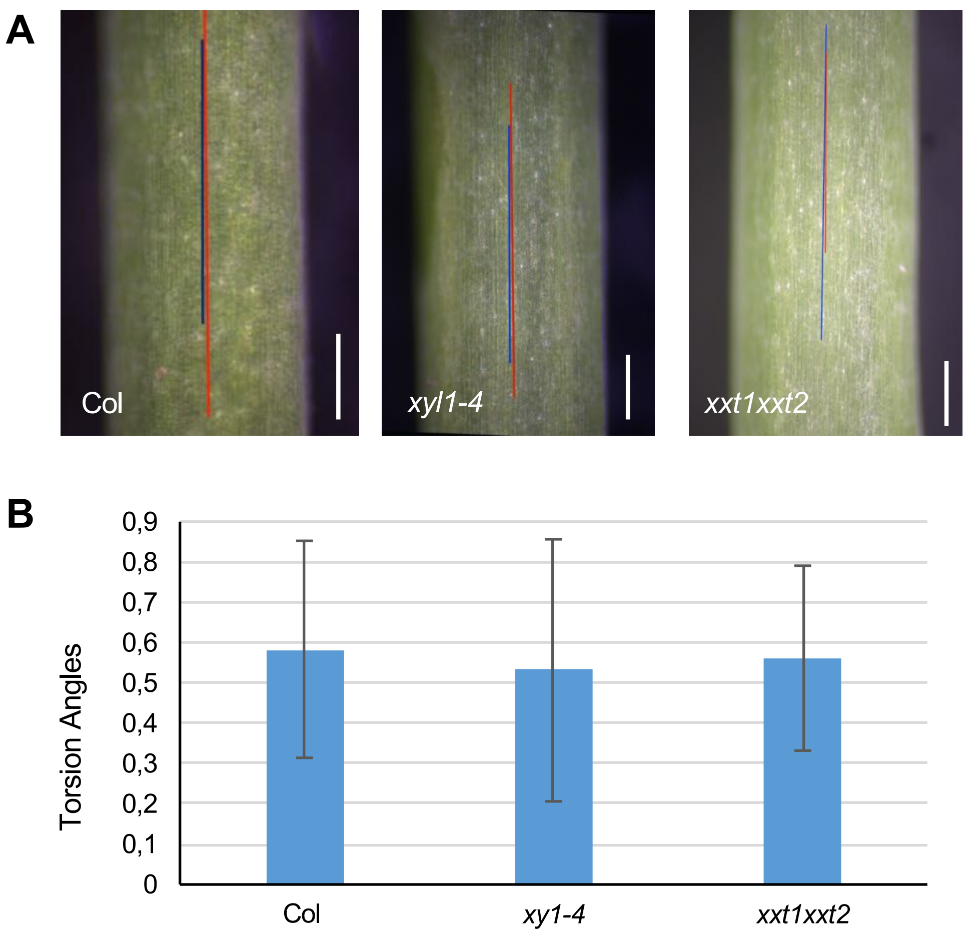


**Figure S4. No apparent torsion on *xyl1-4* and *xxt1xxt2* inflorescence stems.**

(A) Internodes of Col, *xyl1-4* and *xxt1xxt2* stems. Red lines mark the axial direction of the stems; blue lines mark the orientation of cell files on the epidermis.

(B) Quantification of torsion angles of epidermal cells on Col (17 internodes of 3 plants), *xyl1-4* (19 internodes of 4 plants) and *xxt1xxt2* (26 internodes of 6 plants) stems. There are no significant differences among the plants. Scale bars,100µm in (A).

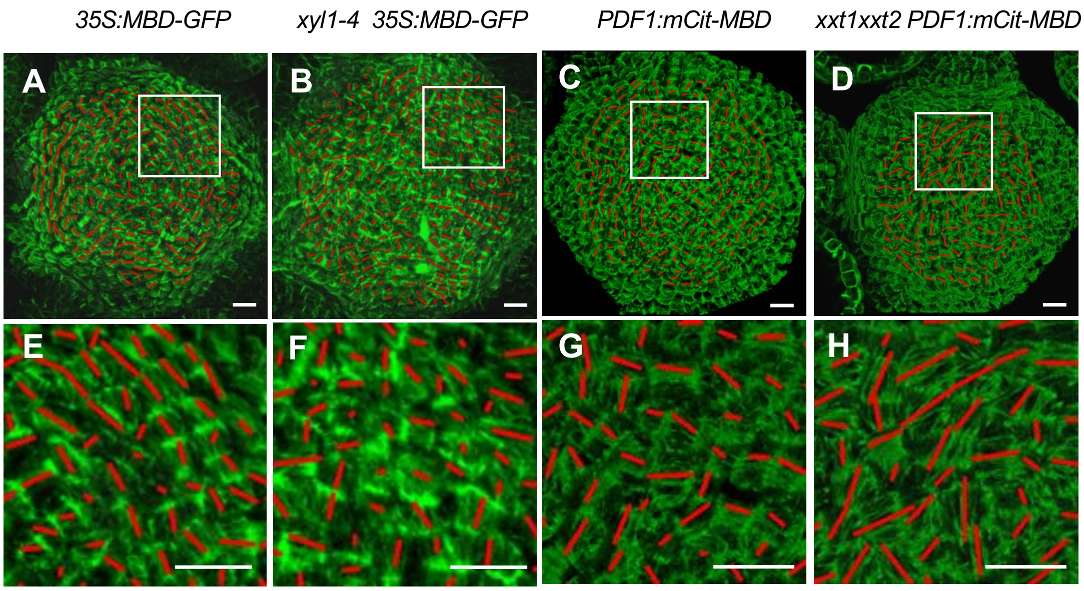


**Figure S5. Microtubule patterning on SAMs of wild-type and XyG mutants.**

(A-D) Overview of wild-type (A, C), *xyl1-4* (B) and *xxt1xxt2* (D) shoot apical meristems.

(E-H) Details enlarged of (A-D). Scale bars, 10μm.
